# FedGMMAT: Federated Generalized Linear Mixed Model Association Tests

**DOI:** 10.1101/2023.10.03.560753

**Authors:** Wentao Li, Han Chen, Xiaoqian Jiang, Arif Harmanci

## Abstract

Increasing genetic and phenotypic data size is critical for understanding the genetic determinants of diseases. Evidently, establishing practical means for collaboration and data sharing among institutions is a fundamental methodological barrier for performing high-powered studies. As the sample sizes become more heterogeneous, complex statistical approaches, such as generalized linear mixed effects models, must be used to correct for confounders that may bias results. On another front, due to the privacy concerns around Protected Health Information (PHI), genetic information is restrictively protected by sharing according to regulations such as Health Insurance Portability and Accountability Act (HIPAA). This limits data sharing among institutions and hampers efforts around executing high-powered collaborative studies. Federated approaches are promising to alleviate the issues around privacy and performance, since sensitive data never leaves the local sites.

Motivated by these, we developed FedGMMAT, a federated genetic association testing tool that utilizes a federated statistical testing approach for efficient association tests that can correct for arbitrary fixed and random effects among different collaborating sites. Genetic data is never shared among collaborating sites, and the intermediate statistics are protected by homomorphic encryption. Using simulated and real datasets, we demonstrate FedGMMAT can achieve the virtually same results as pooled analysis under a privacy-preserving framework with practical resource requirements.

## I. Introduction

With the surge of genomic data generated by Next Generation Sequencing (NGS) [1], [2], [3], [4], number of available genomes have surpassed millions. These data provides opportunities to map genetic factors underlying complex diseases and phenotype. Arguably, Genome-Wide Association Studies (GWAS) are the most popular methods for uncovering the genetic determinants of phenotypic variation [5], [6], [7]. State of the art GWAS tools rely employ mixed modeling to assess the statistical significance between variant genotypes and phenotypes while accounting for the biases in ancestry and cryptic relatedness [8], [9], [10], [11]. Among these methods, linear mixed models (LMMs) have been popularly used for analyzing continuous traits, their intrinsic assumptions (uniform distribution of residuals with respect to variant allele frequency) may bias analyses of the binary traits. For these cases, generalized linear mixed models have been shown to provide unbiased results [12].

Although statistical modeling has advanced in recent years, the real utility of data can be only realized when large sample sizes and more diverse populations are available to identify salient genetic variants. However, it is challenging to build centralized repositories of large datasets mainly due to privacy concerns and data usage agreements[13]. Numerous studies demonstrated that genetic data is very identifying of its owner and their families[14], [15], [16], [17], [18], [19], [20]. Due to very high dimensional nature of genomic data (i.e., very large number of variants), sharing even summary statistics, and other related intermediate data, or simply the existence of variants (e.g., beacons[21], [22], [23]) can create risks of membership inference[24], i.e., whether an individual with known genotypes participated a study. These studies have led to an increased public proclivity against sharing genomic data, related intermediate statistics, and an increased protection of GWAS datasets. Although these restrictions have been partially relieved by NIH in 2018[25], [26], [27], [28], open sharing of raw genotype-phenotype data is still not allowed. In parallel, sharing of data from underserved, historically isolated, and vulnerable populations (Ashkenazi Jewish community, Havasupai Tribe) is challenging since these populations are more vulnerable for stigmatization and further isolation[29], [30], [31]. Due to these concerns, the genetic and biomedical data analysis and research is even more challenging among these communities[32]. Further concerns are related to unauthorized usage of genetic data for solving cold cases[33], usage of genomic data for probing and potential risk of diverting genealogy databases[34].

With the increasing concerns about misusing genome data, regulations like Health Insurance Portability and Accountability Act (HIPAA) and General Data Protection Regulation (GDPR) have been proposed and implemented to protect individual health records including genetic data [35]. The privacy concerns make it untenable to perform, high-powered and high-quality GWAS because these studies require very large sample sizes [36], [37], [38]. As a result, genome data repositories such as UK Biobank [3] often release data with strong restrictions on how data can be used with no possibilities of sharing. Besides the limited data accessibility, many more healthcare data are siloed and underutilized due to concerns around privacy. Some initiatives must be taken to connect these distributed data islands and fully use them while protecting privacy.

Federated Learning (FL) is a privacy-preserving method to bridge isolated data silos without sharing the actual data. Such a property complies with the regulations on PHI protection and provides an alternative to collaborated GWAS across silos. Although some related research has been proposed to provide secure GWAS, however, most of them were focusing on adopting federated methods in *χ*^2^ statistics test [39], [37], Principal Component Analysis (PCA) [40], [41], [42], and linear/logistic regression tests [39], [41]. Among these approaches, there is currently a lack of focus on correcting the confounding by random polygenic effects while performing association testing, mainly due to computational complexity. In this paper, we present a secure federated GWAS algorithm, FedGMMAT, that adopts an efficient and flexible two-step score testing [12], [43], [44]. Our approach integrates homomorphic encryption to protect intermediate statistics, and locally sensitive datasets are never shared among the collaborating entities. We used real and simulated datasets to demonstrate the accuracy and practicality of FedGMMAT in terms of resource requirements.

The source code and example data is available in GitHub repository https://github.com/Li-Wentao/FedGMMAT.

## II. Results

### A. FedGMMAT Algorithm

In the basic setup of FedGMMAT consists of the collaborating sites with local phenotype, genotype, and covariate datasets. The sites use also have access to the design matrices that are used to define the random effects among the individuals, e.g., genetic kinship matrix, which can be computed using a secure kinship estimation method[32]. A central server is used for aggregating intermediate statistics from all sites. Central server is assumed to be a trusted computing entity (no collusions) and is responsible for setting up one pair of asymmetric homomorphic encryption keys (i.e., public, private keys) and sharing the public key with all the collaborating sites[45]. After the keys are setup and public key is shared among all sites, FedGMMAT first executes the null model fitting using a round-robin schedule among the sites wherein each site locally updates the model parameters, encrypts the intermediate results and passes them to the next site to be securely aggregated. The final results are passed to the central server which decrypts the aggregated matrices, updates the global model parameters, and shares them with all sites. It should be noted that null model fitting does not utilize any of the sensitive genotype data. After model parameter convergence, FedGMMAT fits the mixed effect model parameters using a similar round-robin algorithm. The final stage of FedGMMAT is assignment of the score-test statistics to each variant. In this step, central server computes an aggregated projection matrix from all sites. The significance level for each variant are calculated using the score test statistics at each site.

### B. Experimental Setup

We designed 3 different experiments to evaluate our proposed FedGMMAT method.

1. The first experiment is on a synthetic dataset in the baseline model from the R package ‘GMMAT’. This dataset contains 400 samples and 100 SNPs with features of age, sex, and outcome. To mimic the federated learning settings, we randomly split the 400 samples into 3 distinct datasets with sample sizes 124, 120, and 156.
2. The second experiment is on two synthetic genotype datasets that consider population Homogeneity and Heterogeneity. Homogeneous genotype dataset was simulated using the 1000 Genomes Project data using Central Europeans living in Utah (CEU) population by sampling of variants randomly for each subject. For simulating kinship among the subjects, we used a 4-level 16-member pedigree containing 8 founder members and 8 descendants[32]. In the simulation, the founders’ genotypes are sampled, which were used to probabilistically simulate descendant genotypes. We first generated 400 pedigrees (6,400 subjects), which were subsampled down to 6,000 subjects. The overall 6,000×6,000 kinship matrix was estimated using SIGFRIED[32]. For simulating the phenotypes, we selected 20 random causal SNPs and assigned random effect sizes to each from normal distribution *N* (0, 0.5). Environmental effect size was randomly sampled from *N* (0, 0.5) for each individual and gender effect size was set to fixed level of 0.1. Phenotype information was simulated evaluating a logistic link function on the weighted linear combination of covariates and genetic effects on each individual. Population covariates for both homogeneous and heterogeneous datasets were estimated by projection of the genotype data onto 1000 Genomes reference panel [46]. For heterogeneous sample, the genotype data was generated from 3 populations (GBR, YRI, MXL). Both synthetic datasets comprise 6,000 subjects with 62,375 SNPs and 6 covariates (5 population-level PCs and gender). We also preprocessed the genotype data by removing SNPs with minor allele frequency (MAF) less than 0.01, the homogeneous dataset has 56,478 SNPs left and the heterogenous dataset has 62,250 SNPs left. To mimic the federated learning settings, we randomly split the 6,000 samples into 3 sites. Each site has a sample size of 1973, 2037, and 1990 respectively.
3. The third experiment is on 2,545 samples with 571,135 SNPs and 5 covariates (4 PCs and gender). The data is derived from dbGaP (accession identifier phg000049) with Alzheimer’s disease as the phenotype. After filtering out SNPs with MAF less than 0.01, we have 551,062 SNPs. To mimic the federated learning settings, the 2,545 samples are randomly split into 3 distinct datasets with samples size 825, 835, and 885.

### C. Concordance of Variant Association

We first compared the concordance between FedGMMAT and GMMAT under the scenarios described previously.

#### 1) Experiment 1 (400 samples in R package ‘GMMAT’, 100 SNPs)

First, we evaluated the performance of FedGMMAT by using the same synthetic datasets in R package ‘GMMAT’. The synthetic dataset includes 400 samples and 100 SNPs. The baseline model in R will fit a GLMM with age and sex as covariates and disease status as the outcome.

The FedGMMAT first created a privacy-preserving federated network for 3 isolated trainers and trained a federated GLMM model with the 3 split datasets with isolated computation environments. The coefficients of fixed-effects and mixed-effects of GMMAT and FedGMMAT are shown in Table I.

**TABLE I.** COEFFICIENTS of GMMAT and FEDGMMAT in experiment 1.

|  | fixed-effect |  |  | variance component |
| --- | --- | --- | --- | --- |
|  | Intercept | age | sex |  |
| GMMAT | 0.4721 | -0.0068 | -0.0864 | 0.3377 |
| FedGMMAT | 0.4721 | -0.0068 | -0.0864 | 0.3377 |

We observed a very high concordance between the estimates of fixed-effects and mixed-effect coefficients when GMMAT and FedGMMAT are compared. Figure 1 shows the differences of P-values in SNPs’ score test between GMMAT and FedGMMAT with log-scale. The plot shows all SNPs’ p-values of GMMAT and FedGMMAT are virtually identical, laying in the diagonal line.

**Fig. 1.**
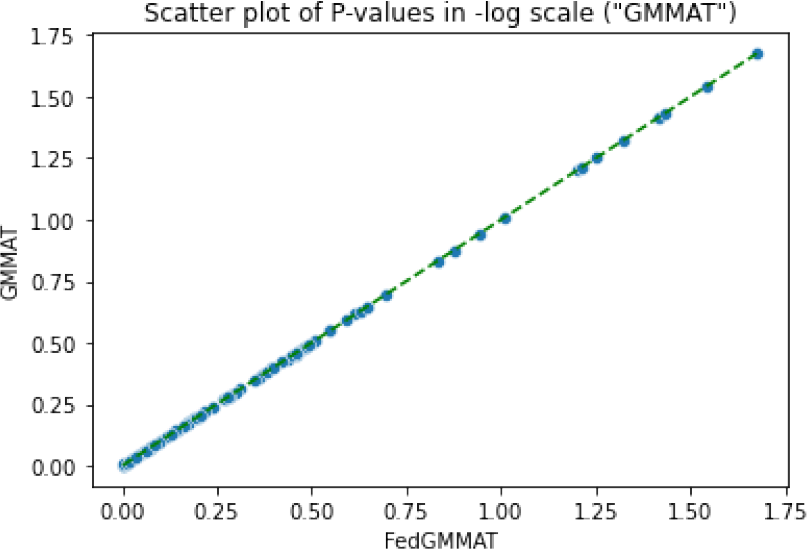
Scatter plot on -log10 scale of P-values of Synthetic data in FedGMMAT and GMMAT.

#### 2) Experiment 2 (6k synthetic samples, 62k SNPs)

Next, we showed the compared the results of FedGMMAT with the synthetic homogeneous and heterogeneous datasets. Similar to the federated settings in Experiment 1, we randomly split the dataset into 3 isolated trainers. Since the homogeneous and heterogeneous settings are sharing the same covariates data, the null models are the same under the two different settings. The model coefficients between GMMAT and FedGMMAT are in Table III. The scatter plot of the p-values in the score test with synthetic is shown in Fig. 2.

**TABLE II.**
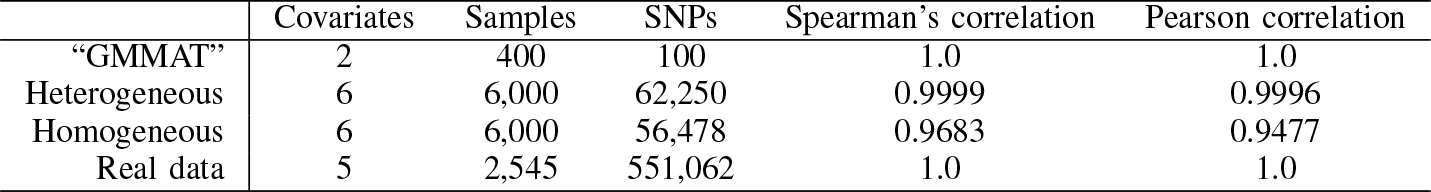
P-values estimation accuracy of FEDGMMAT.

|  | Covariates | Samples | SNPs | Spearman's correlation | Pearson correlation |
| --- | --- | --- | --- | --- | --- |
| "GMMAT" | 2 | 400 | 100 | 1.0 | 1.0 |
| Heterogeneous | 6 | 6,000 | 62,250 | 0.9999 | 0.9996 |
| Homogeneous | 6 | 6,000 | 56,478 | 0.9683 | 0.9477 |
| Real data | 5 | 2,545 | 551,062 | 1.0 | 1.0 |

**TABLE III.** COEFFICIENTS of GMMAT and FEDGMMAT in experiment 2.

|  | fixed-effect |  |  |  |  |  |  | variance component |
| --- | --- | --- | --- | --- | --- | --- | --- | --- |
|  | Intercept | gender | PC1 | PC2 | PC3 | PC4 | PC5 |  |
| GMMAT | -0.0850 | 0.1215 | -0.0011 | -0.0056 | 0.0120 | 0.0238 | -0.0096 | 0.5959 |
| FedGMMAT | -0.0850 | 0.1215 | -0.0011 | -0.0056 | 0.0120 | 0.0238 | -0.0096 | 0.5958 |

**Fig. 2.**
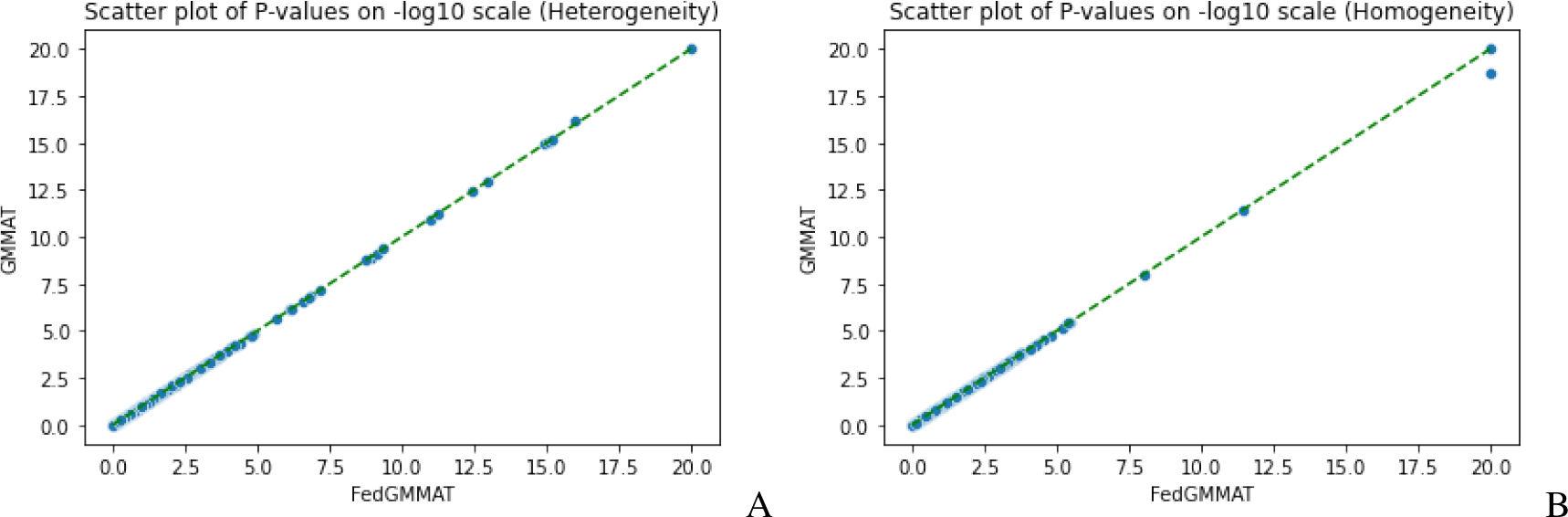
Scatter plot on -log10 scale of P-values in FedGMMAT and GMMAT. (A) Heterogeneity; (B) Homogeneity.

#### 3) Experiment 3 (2.5k dbGaP samples, 551k SNPs)

We finally compared the results from GMMAT and FedGMMAT under the real data usage (dbGaP LOAD Dataset). Among the 3,007 samples with Alzheimer’s disease status, we removed the subjects with unknown phenotype labels, which yields the a dataset with 2,545 subjects who were genotyped at 551,062 SNPs after quality control. In this experiment, we also show the capability of our proposed method with 3 federated trainers. The model coefficients can be seen in Table IV, and the scatter plot of P-values is shown in Fig. 3.

**TABLE IV.** COEFFICIENTS of GMMAT and FEDGMMAT in experiment 3.

|  | fixed-effect |  |  |  |  |  | variance component |
| --- | --- | --- | --- | --- | --- | --- | --- |
|  | Intercept | gender | PC1 | PC2 | PC3 | PC4 |  |
| GMMAT | -0.0038 | 0.0854 | -0.1076 | -0.1316 | -0.0321 | 0.2205 | 0.3997 |
| FedGMMAT | -0.0038 | 0.0854 | -0.1076 | -0.1316 | -0.0321 | 0.2205 | 0.3997 |

**TABLE V.** NOMENCLATURE table.

|  |  |  |  |
| --- | --- | --- | --- |
| $i$ | Index of samples | $\alpha$ | Parameters of covariate |
| $j$ | Index of federated node | $\mathbf{G}_j$ | $n_j$-dim genotype vector of node $j$ |
| $J$ | Total number of federated nodes | $\mathbf{V}$ | $n \times n$ kinship matrix |
| $p$ | Number of covariates | $b_{ij}$ | Random effect of $i$-th sample in $j$-th node |
| $n$ | Total number of samples | $\Sigma$ | $n \times n$ matrix |
| $n_j$ | Number of samples in node $j$ | $\mathbf{b}_j$ | Vector of random effect of $j$-th node |
| $\pi_{ij}$ | Probability of $i$-th sample in $j$-th node | $y_{ij}$ | The outcome of sample $i$ from node $j$ |
| $g(\cdot)$ | Logit link function | $\mathbf{X}_{ij}$ | $p$-dim vector of covariates of sample $i$ from node $j$ |
| $G_{ij}$ | The genotype of $i$-th sample in $j$-th node | $\mathbf{X}_j$ | $n_j \times p$ covariate matrix of node $j$ |
| $\beta$ | Parameters of genotype effect | $\mathbf{W}$ | $n \times n$ diagonal weight matrix |
| $\tau$ | Variance component parameter for random effects | $\Sigma_{(j)}^{-1}$ | $n \times n_j$ matrix of vertically split part of $\Sigma^{-1}$ in node $j$ |
| $\Omega$ | $n \times p$ matrix, product of $\Sigma^{-1}\mathbf{X}$ in node $j$ | $\mathbf{P}$ | $n \times n$ projection matrix |
| $\Omega_j$ | $n \times p$ matrix, product of $\Sigma_{(j)}^{-1}\mathbf{X}_j$ in node $j$ | $\Omega_{(j)}$ | $n_j \times p$ matrix of horizontally split part of $\Sigma^{-1}\mathbf{X}$ in node $j$ |
| $t$ | Index of iteration | | |

**TABLE VI.**
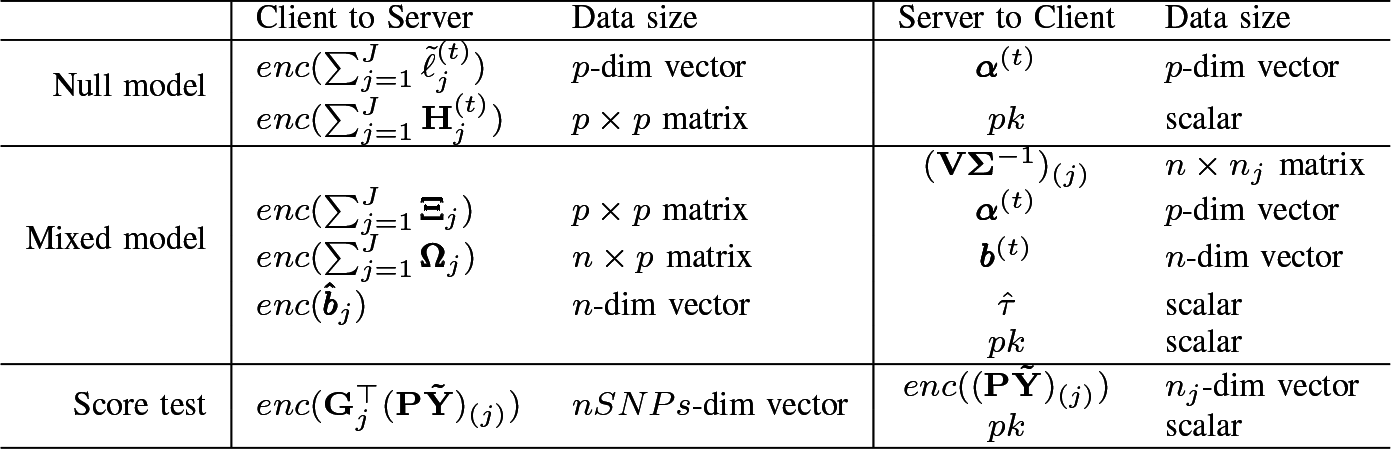
COMMUNICATION information.

**Fig. 3.**
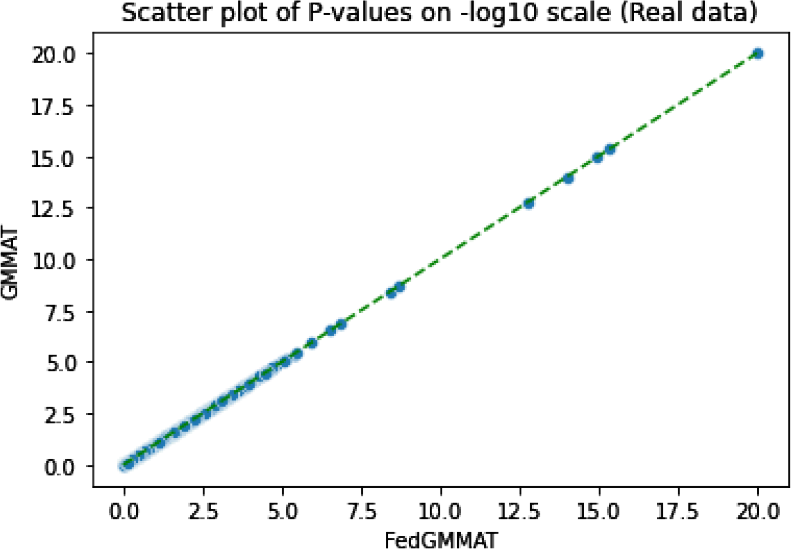
Scatter plot on -log10 scale of P-values in FedGMMAT and GMMAT on real data.

### D. Communication cost performance

Last, we demonstrate the communication cost of our method. We compared the computation time between non-HE protection FedGMMAT and HE protection FedGMMAT. Under the experiment of 6k samples, 62k number of SNPs, and 3-site federated learning simulation, we found the size of the SNPs subset that sequentially feeds in the score test can largely affect the total computation time. The comparison of computation time between HE and non-HE in FedGMMAT can be shown in Fig.4

**Fig. 4.**
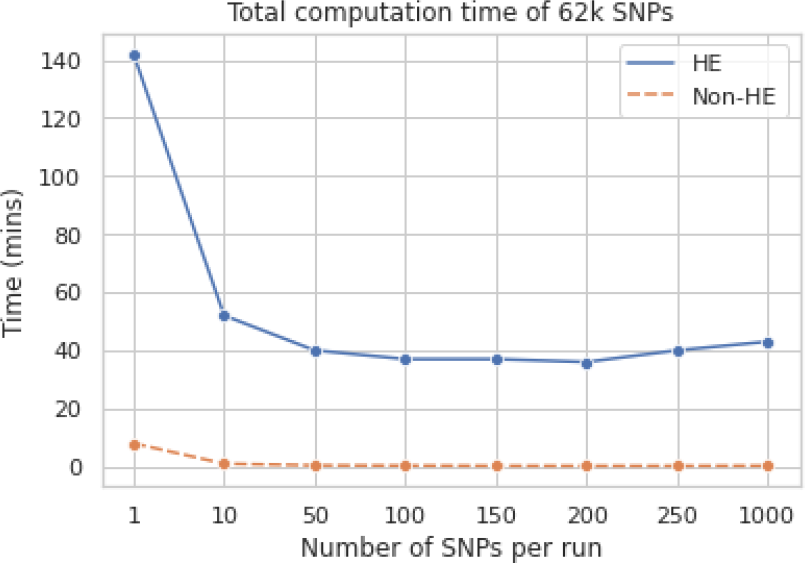
The comparison of computation time with different numbers of SNPs per batch in minutes between HE and Non-HE protection of FedGMMAT.

## III. Discussion

In this paper, we proposed a federated genetic association testing algorithm, FedGMMAT. Comparing the previous method GMMAT, FedGMMAT can bridge the gap of isolated protected data in GWAS and enable international collaborations. FedGMMAT is flexible to integrate multiple subject-level random effects and can be used in different study designs without explicit changes to the algorithmic setup and formulations. This is an inherent advantage that stems from the flexible formulation of the efficient score testing of GMMAT. We also believe that the two-stage significance estimation is advantageous from a privacy preserving perspective because the overall problem is modularized in specific steps of null model fitting and variant scoring where covariates and genotype information is used in different stages. In the future, this can in turn enable a tiered privacy-enabling design for GWAS such that different information is protected at different levels as required by local regulations.

Future work can improve several aspects of FedGMMAT. Currently, the null model information is sent in unprotected form among the sites. However, it should be noted that FedGMMAT’s null model contains very small number of parameters (on the order to 10-20) and we foresee that it is unlikely that the global model parameters can lead to reliable membership or reconstruction attacks.

FedGMMAT assumes an honest-but-curious adversarial model, which is the predominant assumption for the adversaries in genomic data analysis. The central site must be a trusted and non-colluding entity. This can be achieved by setting up a central key management and aggregation service that is operated by a trusted entity (e.g. NIH). Of note, the central server performs lightweight operations and does not bear substantial computational load, which can be implemented as a trusted service. Future studies are necessary to ensure that the malicious entities cannot disrupt the calculations or steal sensitive information.

FedGMMAT currently relies on a round-robin (or cyclic) schedule, which requires sites to wait for their turns without asynchronous updates. In addition, the current implementation relies on TenSEAL framework which may not be optimal for the communication bandwidth. Thus, our future work will focus on refining the current method by adding secure multi-computation protections, embedding a tree-based asynchronous communication framework, and using lightweight communication protocols.

## IV. Methods

### A. Problem setup

We are interested in the single-variant test under federated learning settings; consider the following logistic mixed-effects model:

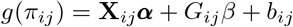

where *g*(·) is the logit link function, and *π*_*ij*_ = *P* (*y*_*ij*_ = 1|**X**_*ij*_, *G*_*ij*_, *b*_*ij*_) is the probability of a binary trait (i.e., disease status) for subject *i* in the federated node *j*. Subscripts *i* and *j* denote the specific subject *i* in the federated node *j*.

**X**_*ij*_ is a *p*-dim vector of covariates, *G*_*ij*_ is a scalar of genotype. When referring to group-level data in federated node *j*, we simply upgrade the subscript to ***X***_*j*_, ***G***_*j*_ and **b**_*j*_. Notice that 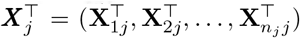 is a *n*_*j*_ × *p* matrix and 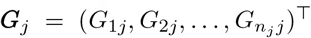 is a *n*_*j*_-dim vector, and 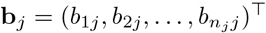 is a *n*_*j*_-dim vector.

For the parameters, ***α*** is a *p*-dim vector that refers to the fixed covariate coefficient, and *β* is the genotype effect. We also assume ***b*** ∼ 𝒩 (0, *τ* **V**) is a *n*-dim vector where 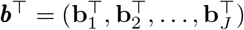, *τ* is the variance component parameter for random effects, and *V* is an *n*×*n* matrix refers to kinship relationship matrix for all samples across the federated nodes, which can be calculated using secure kinship estimation methods[32]. Also define a variance function *v*(·), where *v*(*π*_*ij*_) = *V ar*(*y*_*ij*_|***b***) = *π*_*ij*_(1 − *π*_*ij*_).

FedGMMAT implements the efficient score test that is formulated by GMMAT algorithm[44], which consists of two main steps. First step is null model fitting, which uses only the covariate information without exchanging any sensitive genotype information among sites. Second step is the scoring step, which makes use of the genotype information. Other than the efficiency, we believe that this approach is advantageous from privacy-preservation aspect because it flexibly separates the model fitting and scoring steps, each of which can be federated independently.

### B. Null model and initial estimation

In order to test the significance of the genotype effect, we need to fit a federated logistic mixed model under null hypothesis *H*_0_ : *β* = 0, the null model of single-variant probability on subject *i* in federated node *j* is

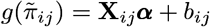

Since the Quasi-Likelihood function (Appendix equation (**??**)) is non-tractable, we use the Laplace method to approximate the integral. Then, we can derive closed-form solutions of fixed-effect coefficients 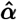 and 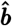 in a federated way. First, we start from getting the initial estimate 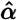 without random effects.

The log-likelihood of the initial estimate of ***α*** under *β* = 0 and *τ* = 0 is of form

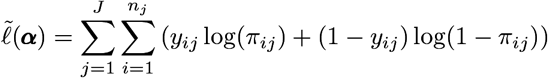

The optimization of such a log-likelihood can be split node-wise; each node can locally calculate its gradient and hessian, then encrypts them to a secure aggregation loop and reconstruct as followings

#### Gradient

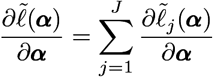

where 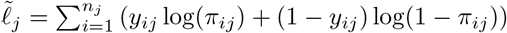

#### Hessian

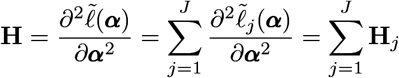

Now the log-likelihood of the null model can be optimized under federated learning settings by Newton’s method

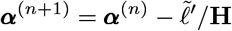

Here, we will derive the optimum 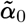 in the null model and denote it as the initial fixed-effects estimates for further computation uses.

### C. Parameters estimation

Denote a local weight vector for site *j*, that is, 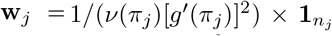, and each node sends it to the center server for construction of global weight matrix 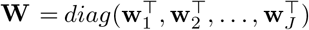. Let **Σ** = **W**^−1^ + *τ* **V** and split it vertically with the sample size of each node, as form 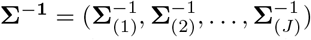.

Also, horizontally split the global matrix 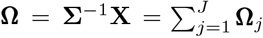 into 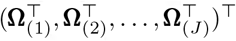. Then send the corresponding part to each node for computation of the following components and send them through the secure aggregation loop

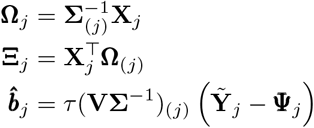

where

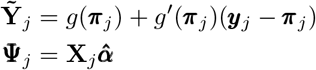

Notice that (**VΣ**^−1^)_(*j*)_ is the *j*-th node vertical split matrix with dimension *n*×*n*_*j*_. And define that (**VΣ**^−1^) = ((**VΣ**^−1^)_(1)_, …, (**VΣ**^−1^)_(*j*)_, …, (**VΣ**^−1^)_(*J*)_). The central server will send the corresponding partition (**VΣ**^−1^)_(*j*)_ to the federated node *j*. Then the federated node will compute and send out the random effect estimate 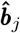. Once the central server receives the components, the fixed-effect coefficients of QL are shown as

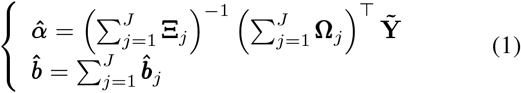

Now we estimate the variance component parameters. We ignore the dependence of **W** on *τ* then adopt Pearson Chi-squared approximation (Equation (**??**)) to the deviance. Then we can optimize the Quasi-Likelihood objective function with respect to variance component parameter *τ* by Average Information REML algorithm [47].

### D. The federated Score Test

Once we derive the optimized estimators, we are ready to conduct the score test on SNPs of interest. First, let us denote a projection matrix **P** with a federated aggregation technique

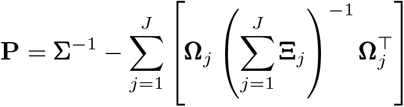

Notice that 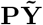 is the residual of the model and calculated in the central server. Then the central server will partition the residual vector 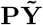 into *J* parts that maches the dimension of local sample size 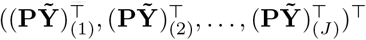 and send them to federated clients accordingly.

Now, the Score test can be shown in the federated settings

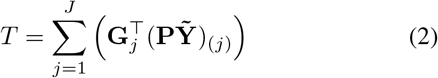

### E. Federated learning workflow and communications

The workflow of the fedGMMAT can be summarized in the following four major steps:

1. **Key Generation and Setup:** The central server generates an asymmetric public-private key pair and shares the public keys with all sites. This step and all homomorphically encrypted operations are implemented using TenSEAL package[45].
2. **Project initiation:** The central server receives the training requests from clients. During this process, model metadata will be collected from each client, including the number of clients in the federated project *J*, the sample size of each client *n*_*j*_, and the number of Covariate.
3. **Federated learning on null model:** In this step, the federated clients will jointly train the null hypothesis model, a logistic regression model, by following steps:
  a. **Broadcast model parameters:** The central server initiates the process of null model learning by broadcasting the parameters and public key to clients.
  b. **Aggregate local information:** Each federated client *j* will calculate and pass the homomorphic encrypted local information (i.e. Gradient 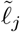 and Hessian matrix **H**_*j*_) with secure aggregation framework in Fig. 6.
  c. **Update model parameters:** The central server will retrieve the encrypted aggregated model information from the last client and decrypt it with the private key (Fig. 6). Then update the model parameters with the aggregated information.
  d. **Broadcast model status:** The central server will first broadcast the up-to-date global parameters to all clients. Then, it will determine whether the model is converged or not by the log-likelihood score. If the difference from the last iteration is within 1×10^−6^, then stop the iteration. If not, goes to step a).
4. **Federated learning on mixed-effects model:** In this step, the federated clients will collaborate on mixed-effect model training. Initializing from the null model,the federated learning model will estimate the fixed-effect coefficients, mixed-effect coefficients, and mixed-effect hyper-parameters with the following steps:
  a. **Broadcast initial model parameters:** The central server will set the parameters from the null model as the initial fix-effect coefficients 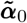, and the variance of targets as the initial mixed-effect hyper-parameters 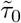, and HE public key. Then send them to the federated clients.
  b. **Aggregate local information:** Once the federated client *j* receives the parameters, it will compute the local information **Ξ**_*j*_, **Ω**_*j*_, 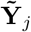, **Ψ**_*j*_ as defined previously. Then cyclically aggregate and send the encrypted aggregated information *enc*(Ξ), *enc*(Ω), 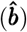 to the central server (Fig. 6).
  c. **Update model parameters:** The central server decrypts the aggregated information then updates variance component estimate 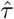, fixed-effect coef-ficient 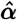, and random effects coefficient 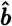.
  d. **Broadcast model status:** After updating and broadcasting the up-to-date model parameters, the central server will determine whether the model is converged with the difference of parameters from the last iteration. If it has not converged, go to step b).
5. **Federated learning on score test:** Once the parameter space 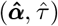 is optimized under the objective function (Equation (**??**)), the FedGMMAT can compute the score test. First, the central server will send the corresponding local model residual 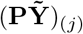 to each federated node. Then the score test statistics *T* can be collected and aggregated from federated nodes by Equation 2.

**Fig. 5.**
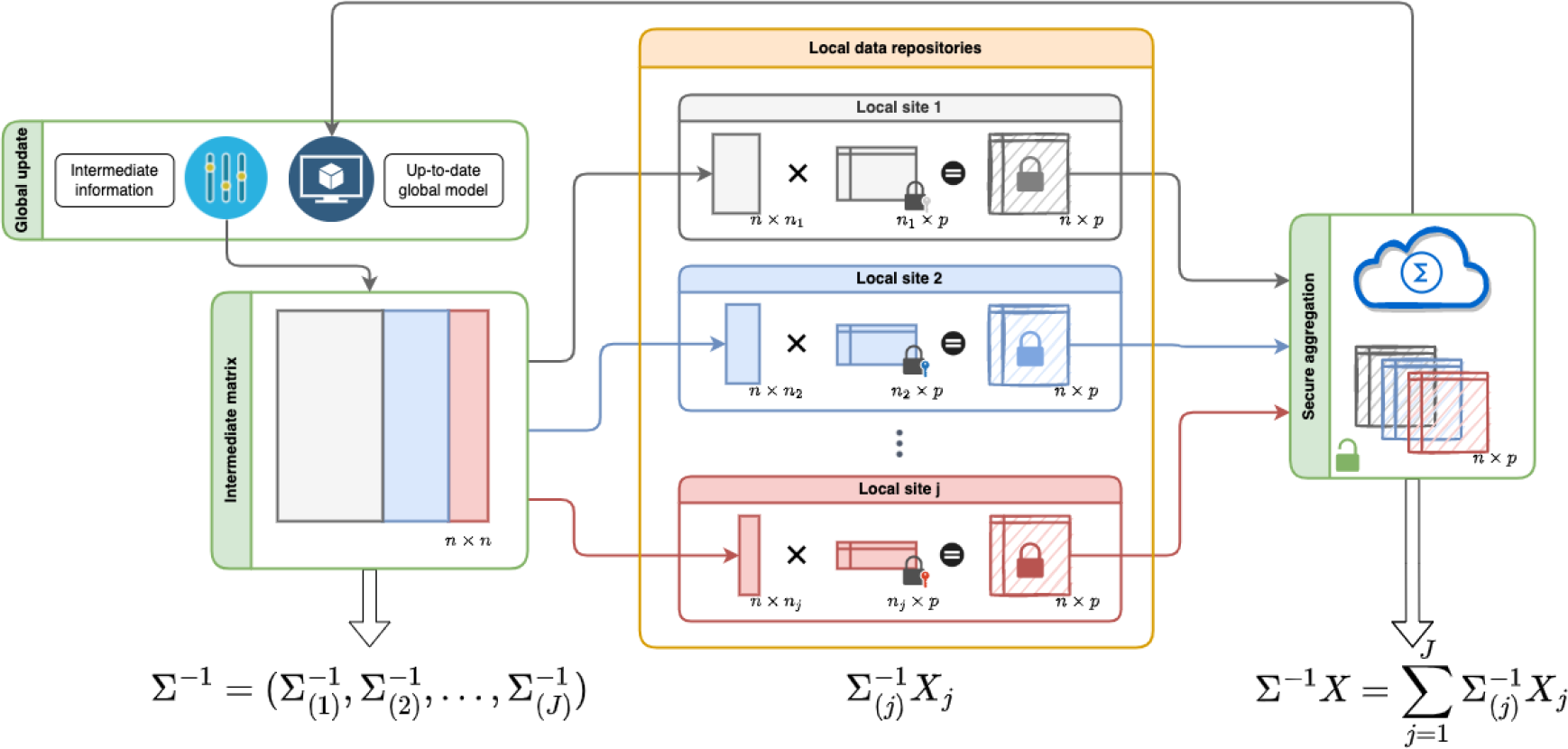
Matrices split and secure computation. In the FedGMMAT framework, each local data repository will maintain its unique dataset locally and gather intermediate model information from Global updates.

**Fig. 6.**
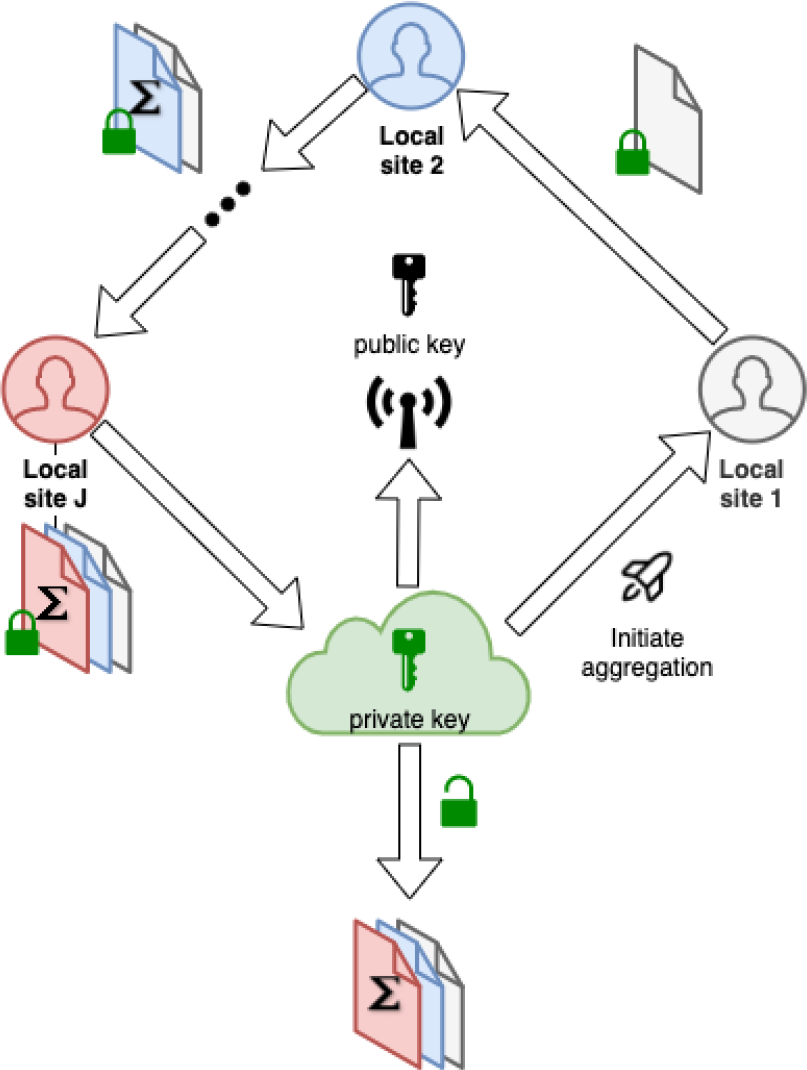
Secure aggregation. During the federated learning training, the aggregation of parameters is protected by Homomorphic encryption through a circle secure aggregation process.

To sum up, the communicated information is shown in Tab. IV, and algorithms 1, 2, 3.

## V. Credit authorship contribution statement

**Wentao Li:** Investigation, Modeling, Method of Study, Data analysis, Original Drafting. **Han Chen:** Review and Editing of the draft. **Xiaoqian jiang:** Review and Editing of the draft, Conception. **Arif Harmanci:** Conception, Data Preprocessing, Review and Editing the draft.

## Acknowledgment

XJ is CPRIT Scholar in Cancer Research (RR180012), and he was supported in part by Christopher Sarofim Family Professorship, UT Stars award, UTHealth startup, the National Institute of Health (NIH) under award number R01AG066749, R01LM013712, and U01TR002062, and the National Science Foundation (NSF) #2124789.

The funders had no role in study design, data collection and analysis, decision to publish, or preparation of the manuscript.

## Algorithm 1: Federated Initial Fixed-effects Estimate

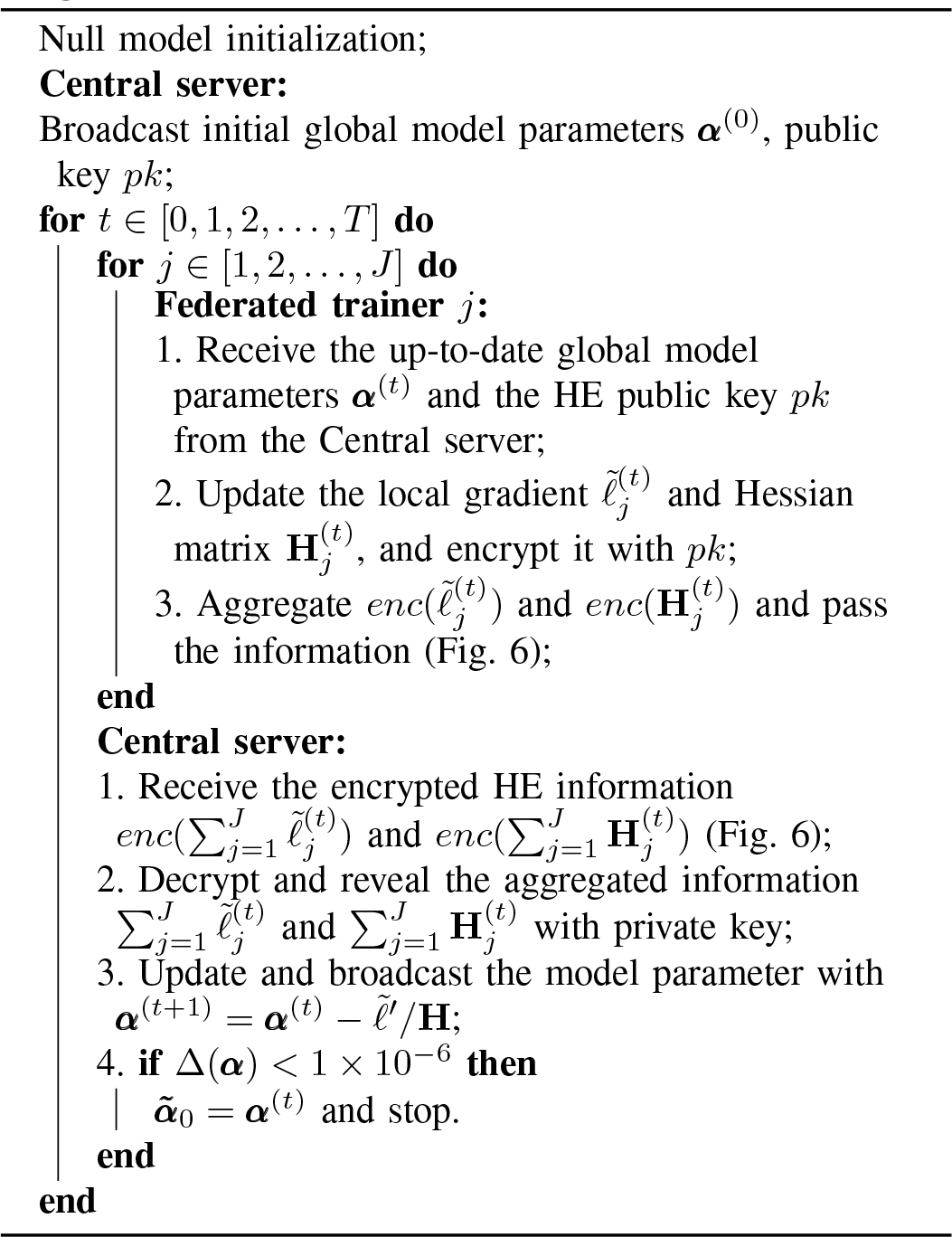

## Algorithm 2: Federated Mixed-effects Model

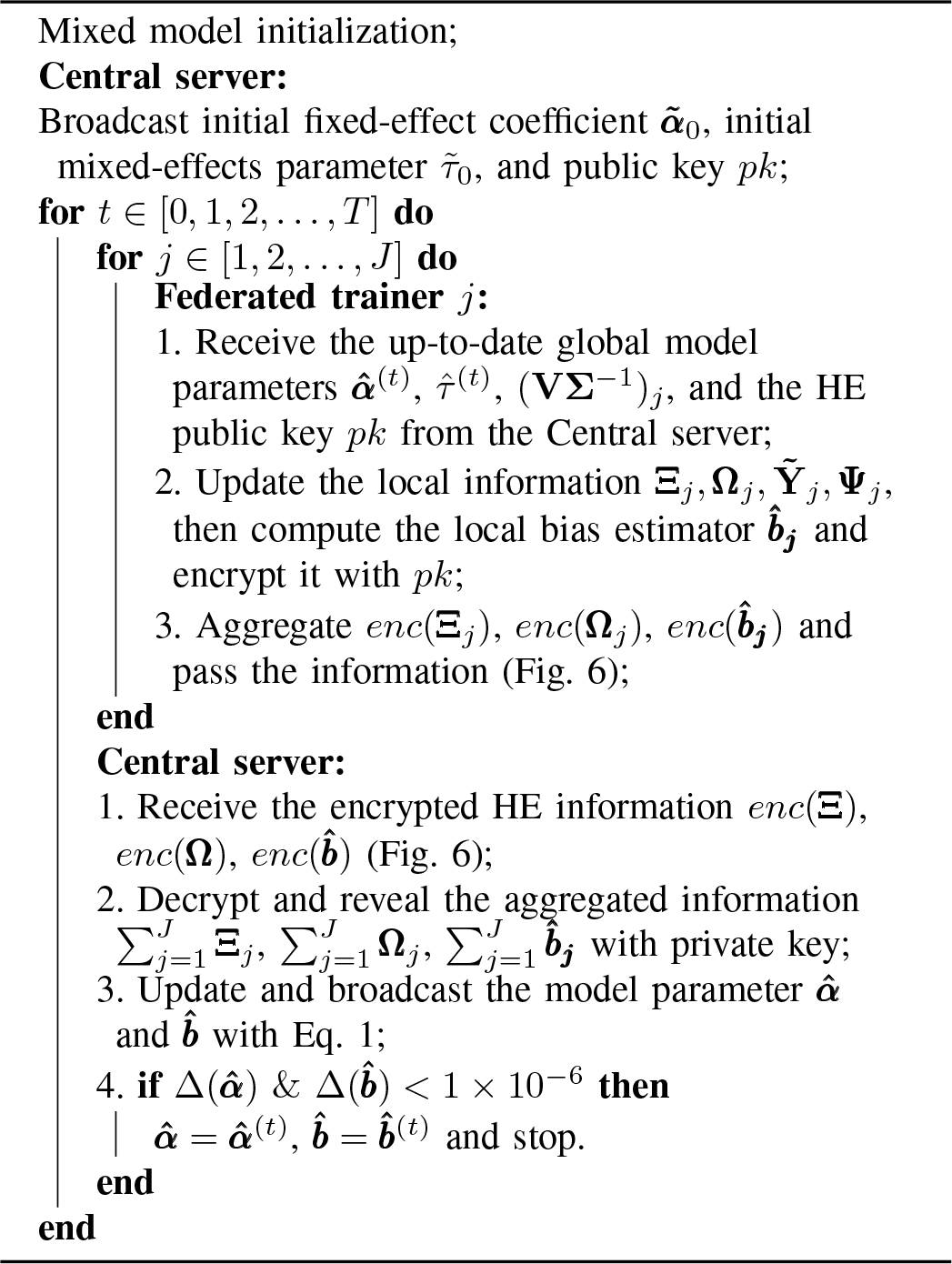

## Algorithm 3: Federated Score Test

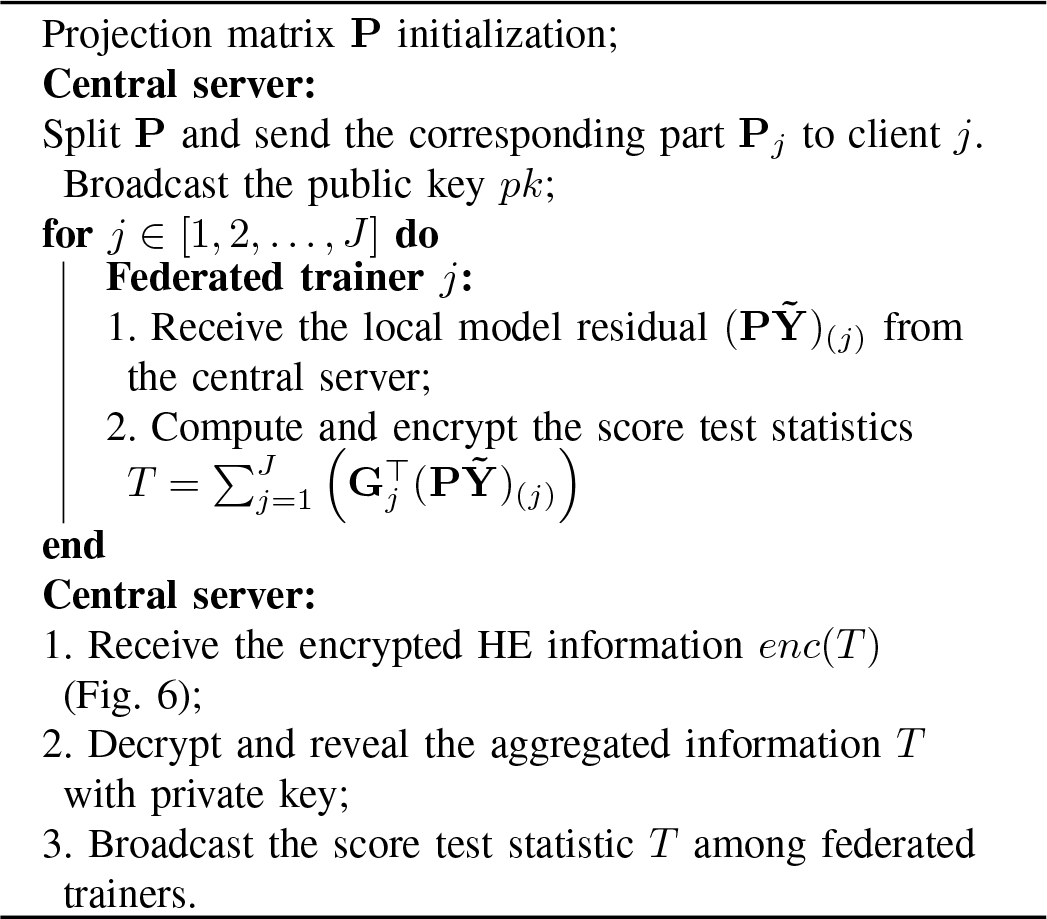

